# A novel crosstalk between Nrf2 and Smad2/3 bridged by two nuanced Keap1 isoforms

**DOI:** 10.1101/2022.11.22.517594

**Authors:** Feilong Chen, Qing Wang, Mei Xiao, Deshuai Lou, Reziyamu Wufur, Shaofan Hu, Zhengwen Zhang, Yeqi Wang, Yiguo Zhang

**Author notes:** Equal contributions to this work (Feilong Chen, Qing Wang).

## Abstract

The Keap1-Nrf2 signalling to transcriptionally regulate antioxidant response element (ARE)-driven target genes has been accepted as key redox-sensitive pathway governing a vast variety of cellular stresses during healthy survival and disease development. Herein, we identified two nuanced isoforms α and β of Keap1, arising from its first and another in-frame translation starting codons, respectively. In identifying those differential expression genes monitored by Keap1α and/or Keap1β, an unusual interaction of Keap1 with Smad2/3 was discovered by parsing transcriptome sequencing, Keap1-interacting protein profiling and relevant immunoprecipitation data. Further examination validated that Smad2/3 enable physical interaction with Keap1, as well as its isoforms α and β, by both EDGETSD and DLG motifs in the linker regions between their MH1 and MH2 domains, such that the stability of Smad2/3 and its transcriptional activity are enhanced with the prolonged half-lives and signalling responses from the cytoplasmic to nuclear compartments. The activation of Smad2/3 by Keap1, Keap1α or Keap1β was likely contributable to a coordinative or another competitive effect of Nrf2, particularly in distinct Keap1-based cellular responses to its cognate growth factor or redox stress. Overall, this discovery presents a novel functional bridge crossing both the Keap1-Nrf2 redox signalling and the TGF-β1-Smad2/3 pathways in healthy growth and development.

## 1. Introduction

Appropriate cell physiology and homeostasis are necessarily maintained by integrating a vast variety of extrinsic and intrinsic signals transducing towards a series of responsive gene expression profiling as so to meet cellular adaptive requirements for changing environments. Of note, Keap1 (Kelch-like ECH-associated protein 1) towards Nrf2 (nuclear factor E2-related factor 2, called NFE2L2)-regulated genes is a key redox signalling pathway, governing transcriptional expression of a big set of antioxidant and electrophile response elements (AREs/EpREs)-driven genes against many of adverse cellular stresses during normal growth and development, and even diseases (1). Within the cells, Keap1 exists in the form of homodimers and interacts with the ETGE and DLG motifs of the Neh2 domain in Nrf2 through its C-terminal Kelch/DGR domain (2,3), enabling Nrf2 to be ubiquitinated by the E3 ligase Cullin-3 for its rapid proteasomal degradation (4). Thereby, it is inferable that under the normal conditions, Keap1 physically binds and sequesters Nrf2 in the cytoplasmic compartments (5,6). Following stimulation by oxidative stress derived from reactive oxygen species (ROS), Nrf2 is released from Keap1 and then translocated into the nucleus, where it is enabled for transcriptional regulation of AREs/EpREs-driven antioxidant, detoxifying and cytoprotective genes. Such target genes include those encoding haem oxygenase 1 (HO-1), NAD(P)H:quinone oxidoreductase-1 (NQO1), γ-glutamyl cysteine ligase catalytic (GCLC) and modified (GCLM) subunits, and glutathione *S*-transferases (GSTs), among others (7-10).

Between the redox-inducible Keap1-Nrf2 pathway and intrinsic developmental signalling cascades, there is an existing key signal-integrated interplay that had been interrogated and explored by a few scientists over the past decade. The intrinsic developmental signalling cascades are predominantly induced by growth factors (including cytokines such as TGF-β (11)), morphogens (e.g., WNT (12)), mitogens and survival factors (e.g., Hedgehog (13)). However, only considerably less attention has been attracted to such a putative as-yet-unidentified interplay between the TGF-β-Smad and Keap1-Nrf2 signalling pathways, even though a large number of studies have indeed elucidated that the transforming growth factor-β (TGF-β) superfamily, also including bone morphogenetic proteins (BMPs), play important roles in the embryogenesis, organogenesis and regulation of the whole-body homeostasis by interacting with membrane receptors that transduce information to the nucleus through both Smad-dependent and-independent pathways (including PI3K-AKT and MAPKs). Notably, the TGF-β receptor-regulated Smad2 and Smad3 (collectively called Smad2/3, hereafter) are two important players of the Smad transcription factor family, that mediates this growth factor-stimulated signalling transduction from the cell membrane surface to the nucleus, and thereby regulates critical physio-pathological functions of target genes involved in healthy growth and development, and even disease development. In this signalling pathway, Smad2/3 serve two downstream receptor-activating proteins of TGF-β1 and binding partners for Smad4 (14). All the Smad family proteins, particularly Smad2/3, are evolutionarily homologous in their structures, with two highly conserved domains, i.e., an N-terminal Mad homology domain-1 (MH1, for binding to target genes) and another C-terminal Mad homology domain-2 (MH2, for binding to cognate partners), joined by a linker region containing four co-phosphorylated serine sites for both GSK-3β (glycogen synthase kinase-3β) and MAPKs (i.e., mitogen-activated protein kinases, including ERKs, JNKs and p38) (15,16). Further studies had also shown that Smad2/3 are closely involved in tumorigenesis, wound healing, immune regulation and the extracellular matrix (17).

Recently, it is elaborately reviewed that many processes that downregulate Nrf2 are *de facto* triggered by TGF-β, with oxidative stress amplifying its signalling (18). The canonical TGF-β pathway leads to Smad2/3- and Smad4-directed increases in the abundances of Hrd1, Bach1, MafK, and ATF3, all of which have been reported to repress Nrf2 activity. Another increase in NOX4*-*mediated production of ROS resulting from Smad2/3 activation by TGF-β heightens two non-canonical pathways *via* activation of TAK1 (TGF-β-activated kinase 1) and TRAF6 (TNF receptor-associated factor 6) signalling cascades to MAPKs, and hence further increases c-Jun levels, which antagonise Nrf2, following dimerization of c-Jun with ATF3 or Fos1/Fra1 (to comprise a functional dimer AP-1). Such oxidative stress may increase suppression of Nrf2 by β-TrCP by facilitating the formation of its DSGIS-containing phosphodegron by GSK-3β together with its priming kinases (e.g., JNK and p38 MAPKs). In turn, it is envisaged that downregulation of Nrf2 by TGF-β can also reinforce activation of both canonical and non-canonical TGF-β signalling pathways if Nrf2-directed antioxidant and detoxification systems are less able to suppress ROS generated by NOX4. Consequently, the increases in Smad-dependent cancer motility and growth, besides intracellular ROS, resulted from Nrf2 deficiency, as accompanied by concomitant phosphorylation of Smad2/3 in their linker regions and C-terminal ends, induction of Slug, a transcriptional repressor of the cell adhesion E-cadherin(19). Conversely, over-expression of wild-type Nrf2, but not its dominant-negative mutant, suppressed the transcriptional activity of a TGF-β1-responsive CAGA-directed luciferase reporter gene, whereas knockdown of Nrf2 enhanced the CAGA reporter activity, as well as the expression of endogenous Smad2/3-target genes, which were further identified to be regulated competitively by a nuclear co-immunoprecipitated complex of Nrf2 with Smad2/3/4 (19). This thus supports the notion that loss of Nrf2 in an oncogenic context-dependent manner also enhances cellular plasticity and motility, at least in part, by the TGF-β1-Smad2/3/4 signalling.

Meanwhile, TGF-β1 is also demonstrated as a potent stimulator of epithelial-to-mesenchymal transition (EMT) by activating the profibrotic genes encoding fibronectin-1 and collagen 1A1 in chronic diseases (20). The TGFβ1-EMT changes are enhanced by *Nrf2* knockdown but suppressed in a stable *Keap1*-knockdown model (to pre-activate Nrf2), concomitantly with repression of TGFβ1-stimulated Smad2/3 phosphorylation and transcriptional activity(20). The Keap1-Nrf2 antioxidant system is further validated as an effective modulator of TGFβ1-stimulated epithelial transition to fibroblastic cells through the Smurf1-Smad7 signalling(20). Consistently, Nrf2 has been shown to inhibit the TGF-β1-dependent expression of fibrosis markers in a human stellate cell line(21,22), whereas TGF-β1 can also reduce the abundance of Nrf2 in another rat hepatic stellate cell line (21,23). However, such inter-inhibitory mechanisms between the Keap1-Nrf2 antioxidant system and TGF-β1-Smad2/3 signalling have not been elucidated to date.

In the present study, we have discovered an unusual interaction of Keap1 with Smad2/3 by parsing transcriptome sequencing, protein profiling, and co-immunoprecipitation data. Further examinations validated that Smad2/3 enable physical interaction with Keap1, including its full-length Keap1α and Keap1β (i.e., Keap1^△1-31^)(24), by both EDGETSD and DLG motifs located in the linker region of Smad2 or Smad3, so that they are segregated in the cytoplasmic compartments, but with their protein stability being enhanced by co-expression of Keap1 or its isoforms. Keap1α and Keap1β were also further demonstrated to monitor the TGF-β1-Smad2/3 signalling transduction of the integral signal from the cell surface to the nucleus. Such activation of Smad2/3 by Keap1, Keap1α or Keap1β was contributable to a putative competitively inhibitory effect of Nrf2, which, in turn, acts on the TGF-β1-Smad2/3 signalling pathways, particularly in distinct Keap1-based cellular responses to its cognate growth factor or redox stress. Overall, this discovery presents a novel functional bridge of the redox-sensor Keap1 with its interactors Smad2/3 involved in growth and development.

## 2. Results

### 2.1 Identification of two Keap1 isoforms α and β

As shown in Figure 1a, two molecular mass-closed protein isoforms of Keap1 were first found to have been differentially expressed in distinct cell lines. Such two electrophoretic bands of Keap1 were determined in HepG2, MHCC97L and THP-1 cell lines, whereas only a single shorter Keap1 protein has been expressed in HL7702, MHCC97H, HCCLM3 and Heap3B cell lines (Fig. 1a). Thereafter, the HepG2 cell line was selectively used for all subsequent studies of two nuanced isoforms of Keap1. Next, distinctive experimental settings were conducted to verify whether its protein expression levels of Keap1 were affected by redox agents. Firstly, total lysates of HepG2 cells were incubated *in vitro* for 1 h with vehicle control (Ctr, distilled water), H_2_O_2_ (0.5 mol/L) or dithiothreitol (DTT, 0.5 mol/L) (Fig. 1b, *left panel*), as previously described (25). Secondly, *in vivo* treatment of HepG2 cells with DTT (1 mmol/L) for different lengths of time (0, 1, 2 or 4 h) and the total cell lysates were then collected (Fig. 1b, *right panel*). Another similar redox treatment of HepG2 cells, that had been transfected with an expression construct for V5-tagged Keap1, was performed (Fig. 1c). These samples were all subjected to Western blotting analysis with distinct primary antibodies. The results revealed that those two protein bands of Keap1 were roughly unaltered in their molecular sizes by DTT and H_2_O_2_ (Figs. 1,b & c, and S1a), no matter whether they were endogenously or ectopically expressed in such distinct redox conditions.

**Figure. 1.**
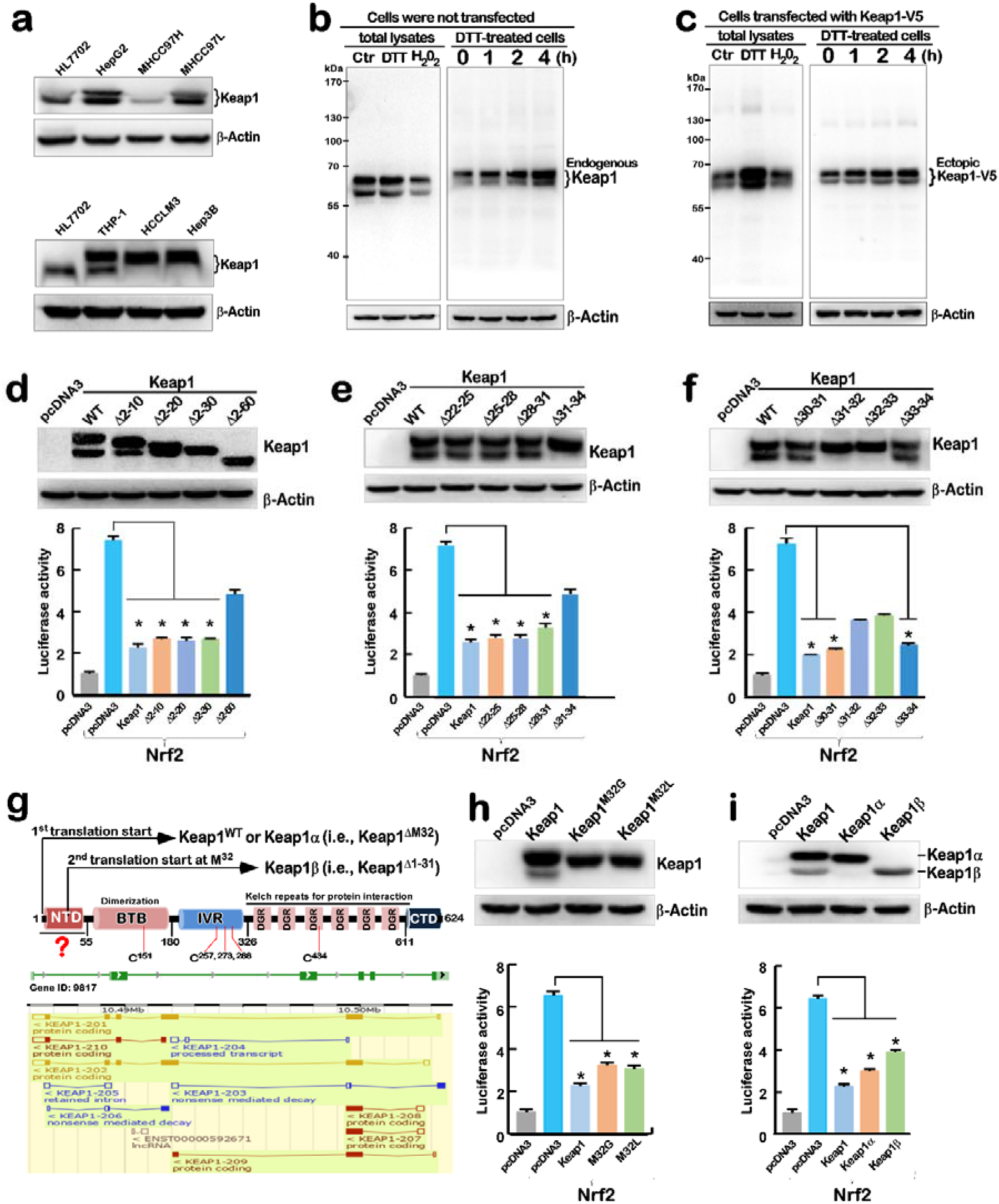
Discovery of Keap1 isoforms α and β. **a.** Differential expression of Keap1 isoforms in distinct cell lines examined by Western blotting with its specific antibody. **b.** The existence of endogenous Keap1 isoforms are unaffected by redox agents. Total lysates of HepG2 cells were treated (*in vitro*) with control water, DTT (0.5 M) or H_2_O_2_ (0.5 M), along with the lysates of HepG2 cells that had or had not been treated (*in vivo*) with 0.1 mM DTT for 0 to 4 h, were separated by SDS-PAGE and visualized by Western blotting with Keap1-specific antibody. **c.** The existence of ectopically-expressing Keap1 isoforms are unaffected by redox agents. Total lysates of HepG2 cells that had been transfected with an expression construct for Keap1-V5 and treated with 0.1 mM DTT for 0 to 4 h, along with the Keap1-expressing lysates being *in vitro* treated with control water, DTT (0.5 M) or H_2_O_2_ (0.5 M), were subjected to Western blotting analysis. **d-f.** Keap1 mutagenesis mapping. HepG2 cells that had been co-transfected with expression constructs for Keap1-V5 and its mutants, along with the Nrf2-expressing plasmid, *ARE-Luc* and *pRL-TK* reporters, were subjected to Western blotting analysis (*upper panels*) and the Dual-Lumi™ luciferase assays as shown graphically (*lower panels*). There data were representative of at least three independent experiments (n = 3 x 3), and significant decreases (*, *p* < 0.01) were determined relative to the positive controls (of Nrf2 with empty pcDNA3 instead of Keap1). **g.** Schematic representation of Keap1 and its isoforms yielded at its mRNA and protein levels. Among them, Keap1α and Keap1β are yielded from alternative translation starting codons at Met^1st^ and Met^32nd^, respectively. All other isoforms of Keap1 were searched from the e!ensembl database (https://asia.ensembl.org/index.html) **h-i.** Identification of Keap1 isoforms α and β. HepG2 cells that had been co-transfected with expression constructs for Keap1-V5 and its mutants (at the Met^32nd^ position), along with the Nrf2-expressing plasmid, *ARE-Luc* and *pRL-TK* reporters, were subjected to Western blotting analysis (*upper panels*) and the Dual-Lumi™ luciferase assays as shown graphically (*lower panels*). These data were representative of at least three independent experiments (n = 3 x 3), and significant decreases (*, *p* < 0.01) were determined relative to the positive controls (of Nrf2 with empty pcDNA3 instead of Keap1).

To further determine distinct lengths of two independent Keap1 isoforms, a series of expression constructs for N-terminally-truncated Keap1 mutants were co-transfected, together with both Nrf2-expressing and *ARE*-driven luciferase plasmids, into HepG2 cells (Fig. 1, d to f). Western blotting results demonstrated that progressive truncation of Keap1’s N-terminal 30 amino acids (aa) enabled its larger isoform (designated Keap1α) to become gradually shortened until this protein disappearance, whereas the mobility of its smaller isoform (called Keap1β) was roughly unaffected by such loss of Keap1’s N-terminal 30-aa (Fig. 1d, *top panel*). By contrast, continuous truncation of Keap1’s N-terminal 60-aa residues (to yield a Keap1^Δ2-60^ mutant) led to the disappearance of Keap1β (besides Keap1α), but it seemed to be replaced by another shorter polypeptide with faster mobility than that of Keap1β (Fig. 2d, *top panel*). This Keap1^Δ2-60^ mutant, rather than others examined, enabled for significant (but not complete) release from inhibition of Nrf2-mediated *ARE-Luc* activity by wild-type Keap1 (Fig. 2d, *lower panel*). Further examination of internal deletions of Keap1 within its N-terminal 34-aa unravelled that the expression of Keap1β, but not Keap1α, was completely prevented by three mutants Keap1^Δ31-34^, Keap1^Δ31-32^ and Keap1^Δ32-33^ (Figs. 1e & 1f, *top panels*), with partial disinhibition of the Nrf2-mediated *ARE-Luc* activity as compared with the case of wild-type Keap1 (Figs. 1e & 1f*, lower panels*).

**Figure. 2.**
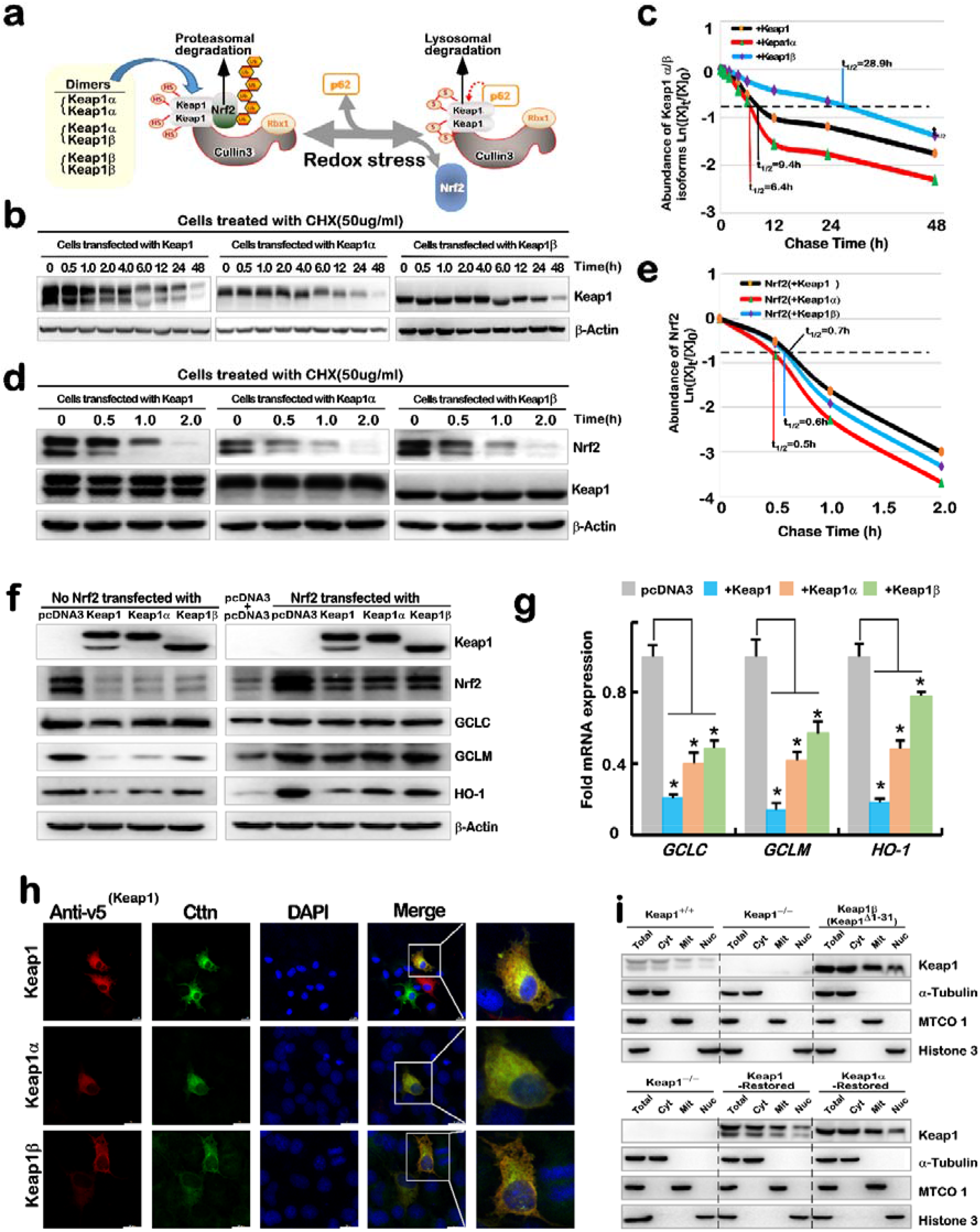
Distinct inhibitory effects of Keap1 and its isoforms (α and β) on Nrf2 stability and *trans*-activity. **a.** Schematic diagram of distinct Keap1 dimers (αα, αβ and ββ) and its interactor Nrf2. In normal (or reductive) conditions, Keap1 interacts with Nrf2 to target this CNC-bZIP factor to the ubiquitin-proteasomal degradation system, but the exposure to oxidative stress, Keap1 disassociates from Nrf2, but instead interacts with p62 targeting to the lysosomal autophagic pathway. **b-e**. Distinct stability of Keap1 and isoforms α and β, with distinct effects on Nrf2 stability. HepG2 cells that had been transfected with expression constructs for Keap1, Keap1α or Keap1β alone (***b***) or plus Nrf2(***d***), were treated with 50 µg/ml of cycloheximide (CHX) for indicated lengths of time in the pulse chase experiments, followed by visualization by Western blotting. The intensity of anti-Keap1 or -Nrf2 immunoblots was quantified by the Quantity One 4.5.2 software (Bio-Rad, CA, USA). The resulting data of Keap1 (***c***) or Nrf2 (***e***) abundances with distinct half-lives were representative of at least three independent experiments, as they are shown graphically, after being calculated by a formula of Ln([A]_t_ /[A]_0_, in which [A]_t_ indicated a fold change (mean ± SD) in each of those examined protein expression levels at different times relative to the corresponding controls measured at 0 h (i.e., [A]_0_). **f**. Distinct effects of ectopic Keap1 Keap1α or Keap1β on Nrf2-targets. HepG2 cells were transfected with expression constructs for Keap1, Keap1α or Keap1β alone (*left panels*) or plus Nrf2 (*left panels*) and then subjected to Western blotting analysis of Nrf2 (including endogenous and exogenous proteins) and its downstream targets. **g**. Differential inhibitory effects of ectopic Keap1, Keap1α or Keap1β on Nrf2-target genes. HepG2 cells were transfected with expression constructs for Keap1, Keap1α or Keap1β alone or plus Nrf2 (Fig. S1b) and then subjected to real-time qPCR analysis of Nrf2 and its target genes. The data were representative of at least three independent experiments (n = 3 x 3), and significant decreases (*, *p* < 0.01) were determined relative to the corresponding controls (of Nrf2 with empty pcDNA3 instead of Keap1). **h**. Subcellular localization of Keap1, Keap1α or Keap1β with CTTN. COS-1 cells were co-transfected with an expression construct for V5-tagged Keap1, Keap1α or Keap1β, together with a CTTN-Flag plasmid, and then subjected to immunocytochemistry with anti-V5 and -Flag and fluorescent-labelled secondary antibody, followed by DNA-staining with DAPI. The representative images were obtained by confocal microscopy. Scale bars =20 μm. **i**. Subcellular fractionation of Keap1, Keap1α or Keap1β from distinct genotypic cell lines. the cytosolic, mitochondria and nuclear fractions were isolated from five examined cell lines, according to the manufacturer’s instruction, and then subjected to Western blotting analysis of Keap1 and its isoforms α and β. Of note, distinct subcellular compartment-specific markers were represented by α-Tubulin, MTCO1 and Histone3, respectively.

Collectively, the above mutagenesis analysis suggests that Keap1α is yielded from its first translation codon (UTG, encoding methionine), while Keap1β is likely translated from its internal UTG-starting codon between its N-terminal aa 31-33 (as illustrated in Fig. 1g). To confirm the latter notion, two-point mutants of Keap1 at the internal translation start codon encoding Met^32^ into glycine or leucine residues (i.e., Keap1^M32G^ or Keap1^M32L^, in Fig. 1h) were made. As expected, the results demonstrated that translational expression of Keap1β was blocked by those two mutants Keap1^M32G^ and Keap1^M32L^ (Fig. 1h, *upper panel*), but almost no obvious changes in Nrf2-mediated *ARE-Luc* activity as compared to that of its wild-type Keap1 (Fig. 1h, *lower panel*). To gain insight into functional distinctions of Keap1α with Keap1β, the Met^32^ residue was deleted to yield a single protein of Keap1α (i.e., Keap1^ΔM32^), while another single Keap1β was allowed for translational expression upon deletion of the N-terminal 31-aa residues of Keap1 (i.e., Keap1^Δ1-31^, Fig. 1g). The results indicated that such two single proteins of Keap1α and Keap1β were enabled for their respective independent expression with the differential ability to inhibit Nrf2-mediated *ARE-Luc* activity (Fig, 1I, *upper and lower panels*).

### 2.2 Distinct effects between Keap1α and Keap1β on Nrf2 and target genes

When unstimulated, Keap1 dimers physically bind to Nrf2 and target this CNC-bZIP protein to the ubiquitin-proteasomal degradation *via* Cullin3 (8,26). After Keap1 is stimulated by oxidative stress, Nrf2 is released from Keap1 and thus translocated into the nucleus before transcriptionally regulating ARE-battery genes, whereas Keap1 is also allowed for another interaction with p62 to be targeted for autophagy-based degradation(27-29). Since being the case, a model is proposed (in Fig. 2a) to provide a possible explanation of three putative dimers of Keap1α and/or Keap1β, as involved in the signalling response to redox stress (4-7),(30).

The stability of Keap1, Keap1α and Keap1β was determined by their half-lives in HepG2 cells, that been transfected with their respective expression constructs and then treated with CHX (50 µg/ml) for 0 to 48 h (Fig. 2b). The pulse-chase experiments revealed distinctive half-lives of Keap1, Keap1α and Keap1β, which were estimated to be 9.4 h, 6.4 h, or 28.9 h, respectively, following CHX treatment of cells (Fig. 2c). Subsequent examination unravelled that endogenous Nrf2 stability was slightly affected by ectopic expression of Keap1, Keap1α or Keap1β (Fig. 2d), because the half-life of this CNC-bZIP protein was evaluated to be 0.7 h, 0.5 h or 0.6 h after CHX treatment of cells expressing Keap1, Keap1α or Keap1β, respectively (Fig. 2e). Further Western blotting analysis showed that Keap1β appeared to enable for modest release of confined Nrf2, as well as its targets GCLC, GCLM and HO-1, to certain extents, when compared to the inhibitory effects of Keap1 and Keap1α (Fig. 2f, *left panels*). Similar results were also obtained from the cells that had been allowed for co-expression of ectopic Nrf2 with Keap1, Keap1α or Keap1β (Fig. 2f, *right panels*). Moreover, real-time quantitative qPCR demonstrated differential inhibition of the mRNA expression levels of Nrf2-target genes *GCLC, GCLM* and *HO-1* by co-expressing Keap1, Keap1α or Keap1β, but with a certain exception of being released by Keap1α or Keap1β, to varying extents (Fig. 2g). Similar effects of Keap1α and Keap1β on ectopic Nrf2-regulated genes were also shown (Fig. S1b).

All the inhibitory effects of Keap1 on Nrf2 and its target genes *GCLC, GSR, GCLM, NQO-1* and *HO-1* were significantly attenuated by knockout of *Keap1^−/−^* (Fig. S1, c & d), whereas Keap1β only retained marginal inhibitory effects on these genes when compared with those controls measured from *Keap1^+/+^* cells. However, their results for Nrf1 were the opposite. Further examination revealed that, when *Keap1-restored* or *Keap1α-restored*, all those inhibitory effects of Keap1 on Nrf2-mediated downstream genes were recovered from *Keap1^−/−^*, but Nrf1 appeared to be upregulated (Fig. S1, e & f). Of note, Keap1-restored cells exerted stronger inhibitory effects than those of Keap1α-restored cells. Such distinction in the inhibitory effects of Keap1, Keap1α or Keap1β on Nrf2 and downstream genes were attributable to their tempospatial co-localization in the cytoplasmic (and even nuclear) compartments. This is supportively evidenced by confocal imaging, illustrating co-localization of V5-tagged Keap1, Keap1α or Keap1β with cortactin (CTTN, an actin-binding cytoskeletal protein, Fig. 2h), as reported previously in the cytoplasm of COS-1 cells (31). Further subcellular fractionation of distinct genotypic cell lines *Keap1^+/+^*, *Keap1^−/−^* and Keap1β (*Keap1^△1-31^*), along with two stably *Keap1-* or *Keap1α*-expressing cell lines ((24)) unravelled that Keap1, Keap1α and Keap1β were recovered primarily in the cytosolic and mitochondrial fractions, and only modestly in nuclear fractions (Figs. 2i & S2). This suggests that they are also likely involved in the nuclear regulatory process, in addition to being responsible for the cytoplasmic and mitochondrial events.

### 2.3 Differential regulation of the TGFβ-Smad2/3 signalling genes and relevant proteins by Keap1α and Keap1β

Distinct genotypic cell lines *Keap1^+/+^*, *Keap1^−/−^*, Keap1β(*Keap1^△1-31^*), *Keap1-restored* and *Keap1α-restored*, that had been confirmed by Western blotting (Fig. S3a) were subjected to transcriptome sequencing to identify differentially regulated gene expression profiles. The results revealed that a total of 1592 differentially expressed genes (DEGs) were common or unique among four examined cell lines, as identified according to the following criteria: fold changes ≥ 2 or ≤ 0.5, plus false discovery rate (FDR) ≤ 0.001 (when compared with equivalent controls measured from *Keap1^+/+^* cells), along with pairwise comparisons of indicated four groups (Fig. S3, b to d), as described previously (24). The top 20 common and differential genes were selected in each of indicated cell lines to map such regulatory targets within their functional association networks (Fig. S3e). Based on the transcriptome data, it is inferable that the expression of most of those genes was consistent, aside from only a few of genes (e.g., *CAMK2D, B4GALNT1, KRT4*, *MAP2*) showed to be expressed in somewhat inconsistent expression trends. This is because 1301 gene products were found to associate with (and even bind to Keap1). For typical example as shown in Fig. 3a, 6 of 13 known Keap1-binding protein-encoded genes (32), which included *MCM3, CHD6, PPP1R13L, AMER1, DPP3* and *PGAM5,* were predominantly expressed in *Keap1^+/+^* cells, except that *PPP1R13L* was partially expressed in Keap1β(*Keap1^△1-31^*) cells. Conversely, *CDKN1A* (i.e., p21 or Cip1, a key regulator of cell cycle) and *SQSTM1* (also known as p62, as a key receptor for autophagy) were highly expressed in *Keap1^−/−^* cells, whereas the other 5 genes *LAMA1, IKBKB, PALB2, NFE2L2*/*Nrf2* and *BRCA2* were dominantly expressed only in *Keap1α-restored* cells.

**Figure. 3.**
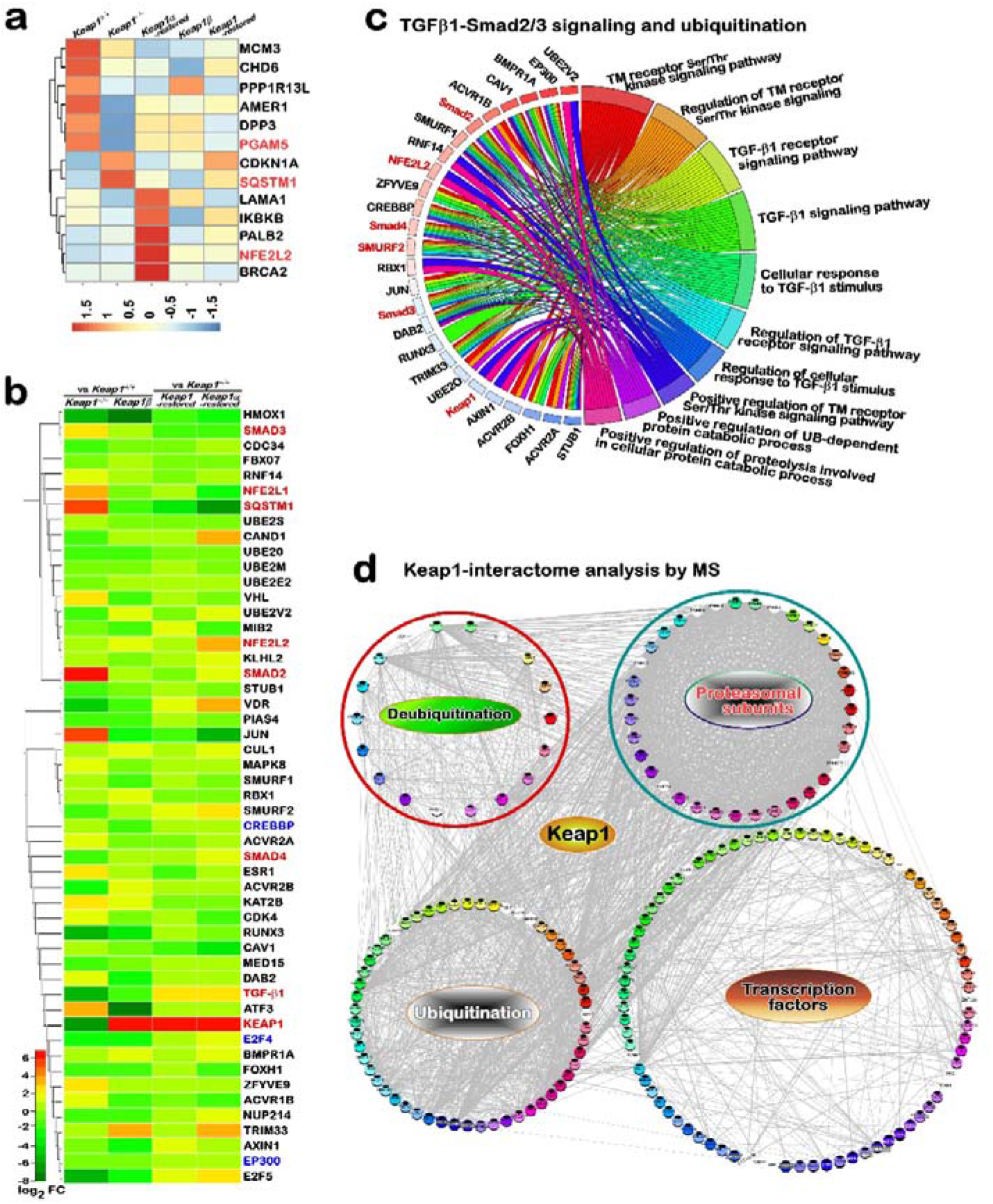
An association of Keap1 with the TGFβ-Smad2/3 signalling discovered by parsing its interactome and transcriptome. **a.** The heapmap shows differential expression levels of those known Keap1-binding protein-encoded genes in distinct cell lines. **b.** The GO enrichment analysis of such genes associated with Keap1, Smad2 and Smad3, which were denoted for the top 10 GO enrichment functions and linked to the same coloured enrichment function. Of note, the TGFβ-Smad2/3 signalling and selective protein modification by ubiquitination or deubuquitination are likely governed by Keap1. **c.** The heatmap with hierarchical clusters of 51 DEGs shared in all four distinct cells lines. As indicated, *Keap1^−/−^* and *Keap1β (Keap1^△1-31^)* were compared with *Keap1^+/+^*, whereas *Keap1-restored* and *Keap1α-restored* were compared with *Keap1^−/−^*. Distinct nodes in the heatmap are represented by the coloured bars showing their values of log2 (fold change). Upregulation was shown in red and yellow, while downregulation was deciphered in green. **d.** The protein-protein interaction networks with ubiquitination, deubiquitination, proteasomal subunits, and those transcription factors, all of which are directly or indirectly bound to keap1. This was obtained from bioinformatics analysis of Keap1-putdown protein mass spectrometry data.

Interestingly, it was, to our surprise, found that Smad2 was presented among the total 1592 DEGs, and also existed as one of the 50 common DEGs in *Keap1^−/−^* and *Keap1α-restored* cell lines (Fig. S3c). According to the BGI gene systemic (Dr Tom) association clustering analysis of Keap1, Smad2 and Smad3 by the top 10 KEGG pathways, it was unveiled that they were dominantly concentrated on the TGF-β1 signalling pathway, ubiquitin-mediated proteolysis, fluid shear stress, the cell cycle and Wnt signalling pathways. By the top 10 of GO enrichment analysis, their biological functions were also primarily concentrated on the TGF-β1 signalling to Smad2/3/4 pathway with related functions (Fig. 3b). As shown in Fig. 3c, approximately 51 DEGs were identified by their association within all four cell lines, and differential expression levels of such genes are also further presented in a hierarchically clustered heat-map. Among such association-clustering DEGs, only 7 genes, including *ZFYVE9, BMPR1A, TRIM33, CUL1, UBE2O, HMOX1/HO-1* and *NFE2L2/Nrf2*, were upregulated or downregulated, with each having similar expression trends, in all four cell lines. Of note, the other 11 genes *SQSTM1/p62, JUN, Smad2/3, VHL, NFE2L1/Nrf1, ACVR1B, BRE2S, UBE2E2, CAND1, SMURF2, ACVR2B,* except *Keap1* were upregulated or downregulated *in Keap1^−/−^* cells, but their opposite results were obtained from *Keap1-restored, Keap1α-restored* and *Keap1β(Keap1^△1-31^)* cell lines (Fig. 3c). Further comparison of the latter three distinct cases revealed that upregulation and/or downregulation of those examined genes by restoration of Keap1 or Keap1α appeared to be largely in similar fashions, whereas 33 of those genes were downregulated by Keap1β (Fig. 3c).

Of striking note, Keap1 was, for the first time, found to directly bind Smad2 by its mass spectrum analysis (Fig. 3d). By analysis of Keap1-putdown polypeptide profiling, Smad2 was identified to be represented by its peptide sequence VETPVLPPVLVPR, whereas Smad3 was also identified by subsequent experiments. In fact, it was shown that Keap1 could be enabled for direct or indirect interactions with 3123 proteins detected here. Among such possibly interacting proteins, besides Smad2/3, other 72 transcription-related factors (e.g., STAT1/3, YY1, FOXK1, CTCF, NKRF, RELA, JDP2 and TFAM), were also put down by V5-tagged Keap1, together with additional 109 proteins that were decomposed of three distinct groups, which comprised 55, 18 and 36 proteins relevant to ubiquitination, deubiquitination and proteasomal subunits, respectively (Fig. 3d). This discovery has unravelled that Keap1 is a crucial multifunctional player for governing and/or maintaining robust proteostasis, because it cannot only contribute to the ubiquitin-mediated proteasomal degradation system by interacting with so many as ubiquitination enzymes (including Cul 1 to 5) along with 26S proteasomal core and regulatory subunits, but also conversely enables for a novel contribution to relevant protein stability by additional deubiquitination system. Furthermore, the results of V5-tagged Keap1α/β-putdown (2914 vs 2972) protein interaction profiling were further subjected to both the GO enrichment analysis and KOG functional annotation (Fig. S4). Keap1α-interacting proteins appeared to be only in the lead of metabolism, particularly for energy production and conversion, protein modification and turnover, and chromatin remodelling. By contrast, Keap1β led to the predominance in multiple aspects, specifically in cellular process and biological regulation by binding organelles or membranes, as well in signalling responses to diverse stimuli from different environments.

### 2.4 Keap1, Keap1α or Keap1β directly interact with two conserved motifs of Smad2/3

Although Smad2 and Smad3 are known to have highly conserved structures and similar biological functions(14-16), it was also, much to our surprise, found by amino acid sequence alignment that two Keap1-binding motifs ETGE and DLG within the Neh2 domain of Nrf2 are represented by additional two homologues EDGETSD and DLQ located within the linker regions of Smad2 and Smad3, respectively (Fig. 4a). By the CABS-dock modelling, it was illustrated that the Keap1-binding peptides could enable it to successfully dock with Smad2 (i.e., YISEDGETSD and NHSLDLQPV) or Smad3 (i.e., YLSEDGETSD and HNNLDLQPV) (Fig. 4b).

**Figure. 4.**
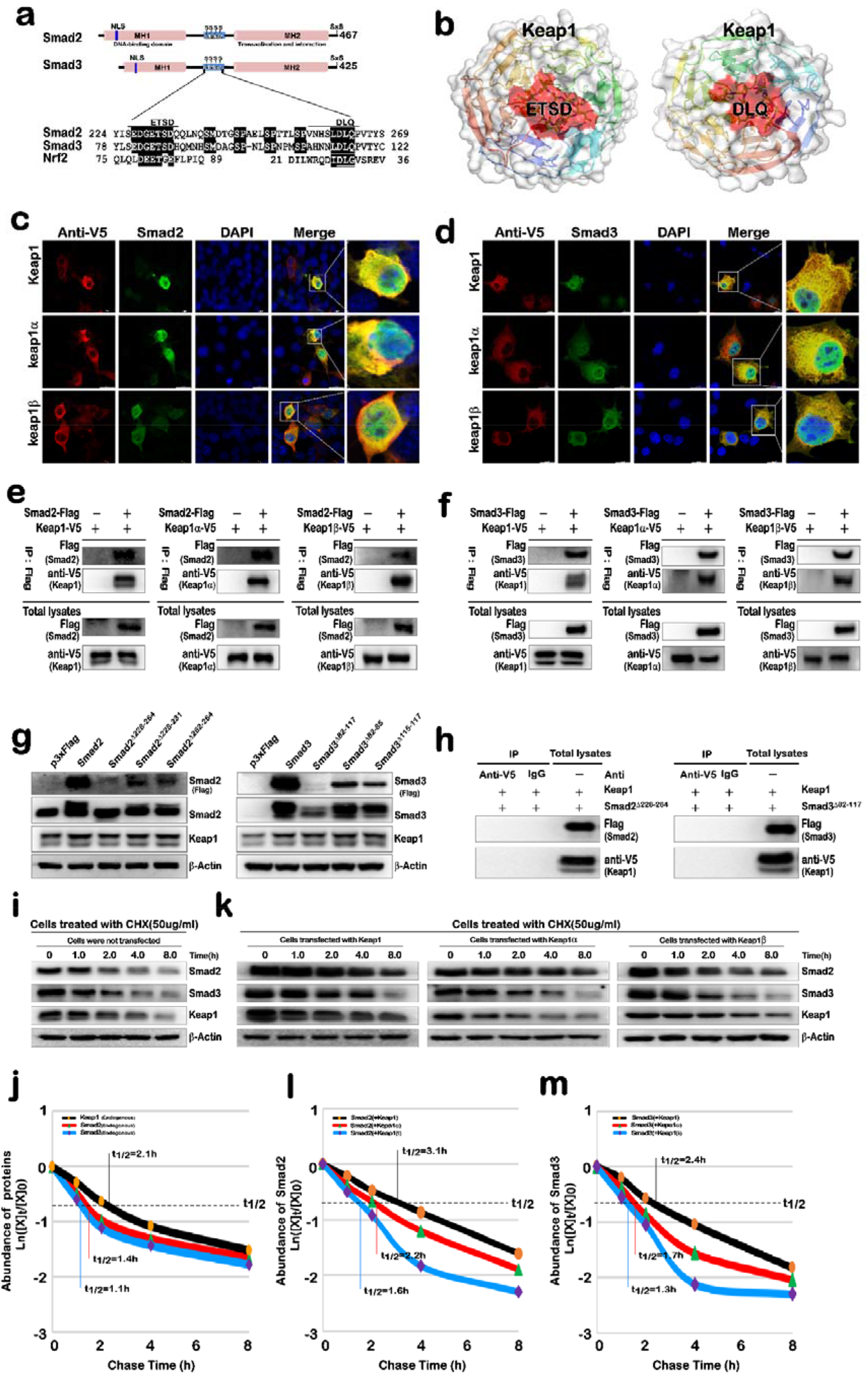
Keap1 and its isoforms α and β interact with Smad2/3 to enhance their protein stability. **a.** Schematic representation of structural domains of Smad2 (NM_005901.6) and Smad3 (NM_005902.4), together with their amino acid sequences within the linker regions being aligned with the known Keap1-binding motifs of Nrf2 (NM_006164.5). **b.** Structural modelling of interaction of Smad2/3 with Keap1 by using the CABS-dock method. The putative Keap1-binding peptides within Smad2 (YISEDGETSD and NHSLDLQPV) or Smad3 (YLSEDGETSD and HNNLDLQPV) were selected for docking into the KEAP1 (PDB code: 1ZGK) with default settings. **c**-**d.** Subcellular co-localization of Smad2/3 with Keap1, Keap1α or Keap1β. COS-1 cells that had been co-transfected with expression constructs for V5-tagged Keap1, Keap1α or Keap1β, together with Smad2-Flag (*c*) or Smad3-Flag (*d*), were subjected to imaging of immunocytochemistry with antibodies against V5 or Flag epitopes, along with the fluorescent secondary antibodies, followed by being stained with DAPI. The immuno-fluorescent images were acquired by confocal microscopy. Scale bars =20 μm **e**-**f**. Co-immunoprecipitation of Smad2 (*e*) or Smad3 (*f*) to Keap1, as well as its isoforms α and β. Total lysates of COS-1 cells that had been co-transfected with expression constructs for Smad2 or Smad3, along with V5-tagged Keap1, Keap1α or Keap1β, were subjected to co-immunoprecipitation with (herein) and antibody (Fig. S8), followed by Western blotting with anti-V5 or -Flag antibodies, respectively. **e.** No effects of Smad2/3 mutants on the abundance of Keap1. Total lysates of HepG2 cells were transfected with expression constructs for Smad2/3-Flag or their mutants and then subjected to determination by Western blotting with antibodies against Keap1, Smad2/3 and their C-terminal Flags. **h**. No immunoprecipitates of Smad2^△228-264^-Flag or Smad3*^△^*^82-117^-Flag were put down by V5-tagged Keap1. Total lysates of COS-1 cells that had been co-transfected with expression constructs for V5-tagged Keap1 plus either Smad2^△228-264^-Flag or Smad3*^△^*^82-117^-Flag, were subjected to co-immunoprecipitation by V5-tagg antibody, followed by Western blotting with Flag or V5 antibodies. **i**-J. The CHX-pulse chase experiments to determine the half-lives of endogenous Smad2/3 and Keap1 expressed in HepG2 cells that had been treated with 50 g/mL CHX for 0 to 8 h, followed by visualization by Western blotting with their specific antibodies. Then the intensity of the immunoblotts representing Keap1, Smad2 or Smad3 was quantified by the Quantity One 4.5.2 software (Bio-Rad, Hercules, CA, USA). The data were representative of at least three independent experiments, and are shown graphically, after being calculated by a formula of Ln([A]_t_ /[A]_0_, in which [A]_t_ indicated a fold change (mean ± SD) in each of those examined protein expression levels at different times relative to the corresponding controls measured at 0 h (i.e., [A]_0_). **k-m**. Differential enhancement of Smad2/3 stability by Keap1, Keap1α and Keap1β. HepG2 cells were transfected with expression constructs for Keap1, Keap1α and Keap1β and treated with 50 g/mL CHX for 0 to 8 h (*k*), and then subjected to the pulse chase experimental analysis of endogenous Smad2/3 stability, which was estimated by their half-lives of Smad2 (*l*) and Smad3 (*m*). The data were representative of at least three independent experiments.

Confocal imaging of immunocytochemistry showed that the red fluorescent signals representing V5-tagged Keap1, Keap1α or Keap1β were superposed with flag-Smad2/3 exhibiting green fluorescent signals primarily in the cytoplasm, but modestly superposed images of Keap1β with Smad2 were also presented in the nucleus (Figs. 4c, 4d). The putative interaction of Keap1, Keap1α or Keap1β with Smad2/3 was further corroborated by a series of co-immunoprecipitation assays, revealing that flag-Smad2/3 enabled for immunoprecipitation of V5-tagged Keap1, Keap1α or Keap1β (Figs. 4e, 4f) and *vice versa* (i.e., V5-tagged Keap1, Keap1α or Keap1β also enabled Smad2 or Smad3 to put down, Fig. S5).

Further mutagenesis mapping of putative Keap1-binding EDGETSD and DLQ motifs in the linker region of Smad2/3 revealed that the resulting mutants caused their protein expression levels to be decreased (Fig. 4g), particularly in the cases of Smad2^△228-264^ and Smad3^△82-117^. The latter two mutants lacked both the EDGETSD and DLQ motifs of Smad2/3 so that their immunoprecipitates were not put down by V5-tagged Keap1 (Fig. 4h), although the expression abundance of Keap1 was unaffected or even slightly enhanced (Fig. 4g). Collectively, these suggest that the stability of Smad2/3 may not only be modulated by their EDGETSD/DLQ-encompassing linker regions, and also monitored by their interacting Keap1, but no obvious differences were made by Keap1α or Keap1β in this process. Of note, the endogenous Keap1 protein (consisting of both α and β isoforms) also enabled to directly immunoprecipitate with endogenous Smad2 or Smad3 in additional two cell lines of MHCC97L and THP-1 (Fig. S6).

### 2.5 Distinct impacts of Keap1, Keap1α and Keap1β on Smad2/3 stability and target gene expression

To further verify whether the protein stability of Smad2/3 was influenced by Keap1, Keap1α or Keap1β, a series of the CHX pulse-chase experiments were carried out (Figs. 4i to 4m and S9). Firstly, the stability of endogenous Smad2/3 was determined by measuring their half-lives which were estimated to be 1.4 h or 1.1 h, respectively, after CHX treatment of cells (Figs. 4i, 4j), while the half-life was evaluated to be 2.1 h (Fig. 4i, 4j). Secondly, it is very interesting to note that forced expression of Keap1 Keap1α or Keap1β caused the endogenous Smad2/3 half-lives to be, to different extents, prolonged (Fig. 4k). As it is indeed, the half-life of endogenous Smad2 was prolonged respectively by ectopic Keap1 to 3.1 h, Keap1α to 2.2 h, and Keap1β to 1.6 h (Fig. 4l), while the endogenous Smad3 half-life was also prolonged by ectopic Keap1 to 3.4 h, Keap1α to 1.7 h, and Keap1β to 1.3 h, respectively, after CHX treatment (Fig. 4m). Thirdly, the exogenous Smad2/3 half-lives were also prolonged by ectopic Keap1 and its isoforms α and β to be much longer than those of their endogenous proteins (Fig. S7). Just so ectopic Smad2 half-life was extended by over-expressing Keap1, Keap1α or Keap1β to reach 4.3 h, 3.1 h, or 2.0 h, respectively (Fig. S7, a & b), whereas ectopic Smad3 half-life was also extended by over-expressing Keap1, Keap1α or Keap1β to reach 4.0 h, 2.7 h or 1.7 h, respectively, after treatment of cells with CHX (Fig. S7, c & d). Altogether, these demonstrate that the stability of Smad2/3 is enhanced distinctively by Keap1, as well as its isoforms α and β, because the latter three can differentially promote Smad2’s half-life more strongly than that of Smad3.

Next, distinct Keap1-based genotypic cell lines were employed to explore the effects of Keap1, Keap1α or Keap1β on the basal expression of Smad2/3-target genes. When compared with *Keap1^+/+^* cells, Smad2, Smad3, Smad4, TGF-β1 and E2F4 (E2F transcription factor 4), PAI-1 (*<u>p</u>*lasminogen *<u>a</u>*ctivator *<u>i</u>*nhibitor 1, also called SERPINE1, a member of the *<u>ser</u>*ine *<u>p</u>*roteinase *<u>in</u>*hibitor) superfamily) and TMEPAI (*<u>t</u>*rans*m*embran*<u>e</u> <u>p</u>*rotein *<u>a</u>*ndrogen induced 1), of which the latter five had been identified as direct downstream target genes of Smad2/3(33-35) were down-expressed in *Keap1^−/−^* cells, but conversely, they were all up-expressed in Keap1β(*Keap1^△1-31^*) cells (Fig. 5a). Similarly downregulated or upregulated changes in their mRNA expression levels of these examined genes were observed in *Keap1^−/−^* or Keap1β(*Keap1^△1-31^*) cell lines, respectively (Fig. 5b). Upon the restoration of Keap1 or Keap1α, all those examined protein abundances (Fig. 5c) and their mRNA expression levels (Fig. 5d) were strikingly promoted, to different extents, in *Keap1-restored* and *Keap1α-restored* cell lines. These results also demonstrated the function of restored Keap1 appeared to be significantly stronger than that of its α-isoform. Further *E2F4-Luc* assays revealed that its transcriptional expression was markedly upregulated by Smad2 or Smad3, and also manifested with the highest expression levels being enhanced by Keap1 and the lowest expression levels elicited by Keap1β (Fig. 5e).

**Figure 5.**
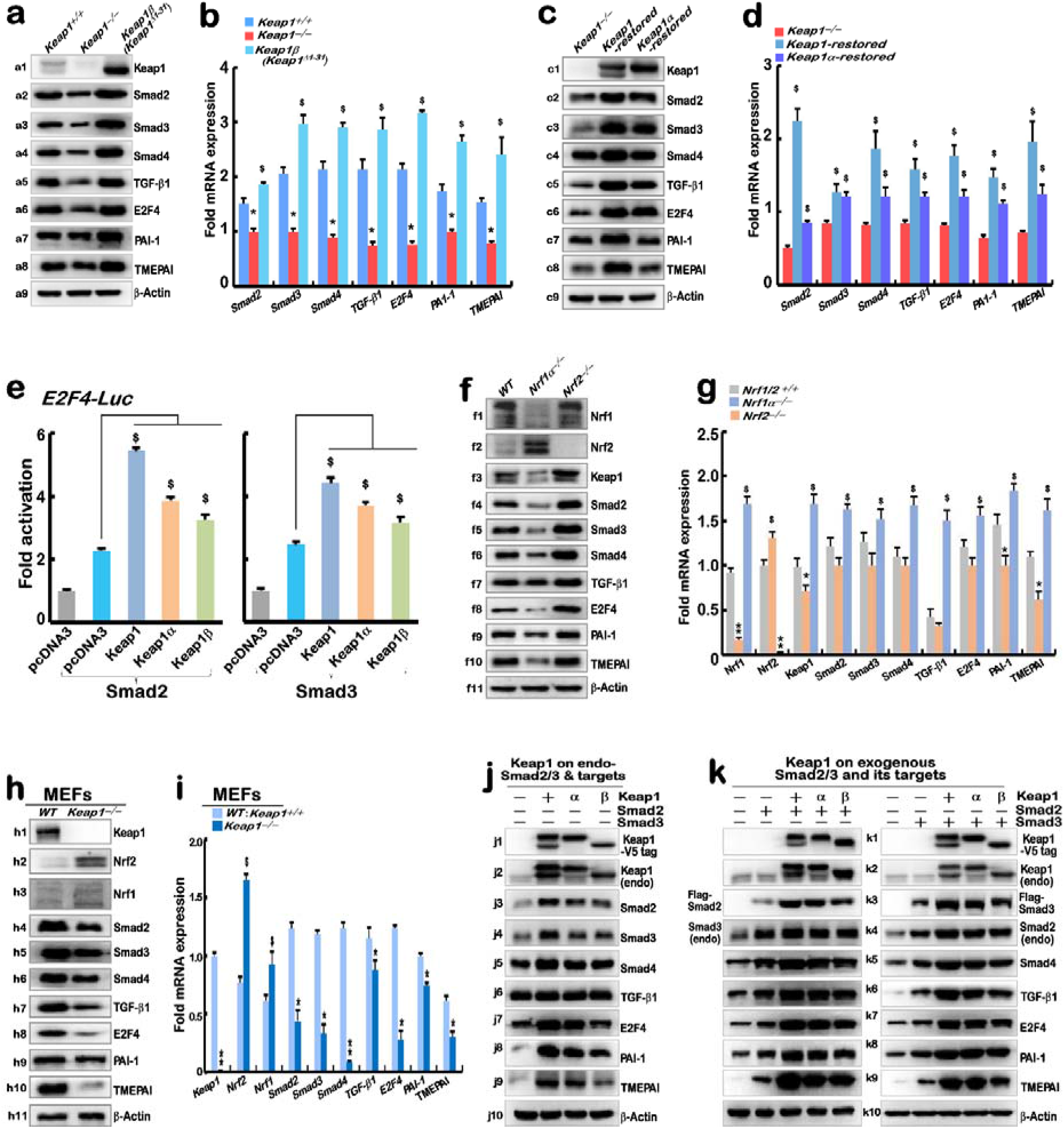
Upregulation of Smad2/3-mediated genes by Keap1 and its isoforms α and β. **a-d**. Distinct effects of Keap1, Keap1α and Keap1β on Smad2/3 and related cognate genes. These protein and mRNA expression levels in *Keap1^+/+^*, *Keap1^−/−^* and *Keap1β(Keap1^△1-31^)* cell lines were determined by Western blotting (*a*) and quantitative real-time PCR (*b*). Similar experiments were carried out to determine the relevant protein and mRNA abundances (*c* and *d*) in *Keap1^−/−^*, *Keap1-restored* and *Keap1α-restored* cell lines. The data were representative of at least three independent experiments (n = 3 x 3), and significant decreases (*, *p* < 0.01) and significant increases ($, *p* < 0.01) were determined relative to the corresponding controls. **e.** The effects of Keap1 and its isoforms (α and β) on Smad2/3-mediated reporter activity. HepG2 cells that had been co-transfected with expression constructs for Smad2 or Smad3, plus Keap1, Keap1α, Keap1β or empty pcDNA3 vector, together with the *E2F4-Luc* reporter gene and *pRL-TK* control, were subjected to the Dual-Lumi™ reporter assay. The Smad2/3-mediated reporter activity was calculated from at least three independent experiments (n = 3 x 3). Significant increases ($, *p* < 0.01) were also determined relative to the corresponding controls. **f**-i Regulation of Smad2/3 and cognate genes by altered Keap1 in human *Nrf2^−/−^* and *Nrf1*α*^−/−^* hepatoma cell lines, and *Keap1^−/−^* mouse embryonic fibroblasts (MEFs), when compared to their corresponding wild-type cell lines. Both the protein and mRNA expression levels of Keap1, Smad2/3 and cognate target genes in the examined three cell lines were determined by Western blotting with each specific antibody (*f, h*) and quantitative real-time PCR (*g, i*). The resulting data were representative of at least three independent experiments (n = 3 x 3), and significant decreases (*, *p* < 0.01) and significant increases ($, *p* < 0.01) were determined relative to the corresponding controls. J-k. Effects of ectopic Keap1 and its isoforms (α and β) on Smad2/3 and target gene products. Each of expression constructs for Keap1, Keap1ααand Keap1β alone (*j*) or plus Smad2 or Smad3 plasmids (*k*) were transfected into HepG2 cells, respectively. After 24 h, the total proteins were collected and subjected to Western blotting analysis of all indicated protein expressions. The empty pcDNA3 vector (−) was also transfected as an internal control. This is a representative of at least three independent experiments (n = 3)

To gain insight into the effects of endogenous Keap1 on Smad2/3, both cell lines of *Nrf1α^−/−^* (with diminished Keap1 abundance(36)) and *Nrf2^−/−^* (with enhanced Keap1 expression) were employed here. As excepted, the results unravelled that Keap1, Smad2, Smad3, Smad4, TGF-β1, E2F4, PAI-1, and TMEPAI were all decreased at their protein abundances in *Nrf1α^−/−^* cells (Fig. 5f), despite only less or no changes in most of their mRNA expression levels except for *E2F4, PAI-1* and *TMEPAI* that were obviously downregulated by such loss of *Nrf1α^−/−^* (Fig. 5g). By contrast, they were all upregulated probably due to Keap1 activation by loss of *Nrf2^−/−^* (Fig. 5, f & g). Collectively, these demonstrate that Keap1 contributes to the positive regulation of Smad2/3 and target genes. This notion is also supported by further experimental evidence, showing the genomic loss of *Keap1^−/−^* led to significant downregulation of Smad2, Smad3, Smad4, TGF-β1, E2F4, PAI-1, and TMEPAI, although Nrf2 was markedly upregulated, whereas Nrf1 was only marginally enhanced, in *Keap1^−/−^* MEFs (mouse embryonic fibroblasts) (Fig. 5, h & i).

Lastly, to further determine the positive effect of Keap1 or its isoforms (α and β) on endogenous and exogenous Smad2/3, wild-type HepG2 cells were co-transfected with expression constructs for Keap1, Keap1α or Keap1β (Fig. 5j), or together with Smad2 or Smad3 (Fig. 5k). As anticipated, the results revealed that distinct extents of increases in the exogenous and endogenous protein expression levels of Smad2, Smad3, Smad4, TGF-β1, E2F4, PAI-1, and TMEPAI were promoted by Keap1, Keap1α or Keap1β (Fig. 5, j & k). Their promotion by Keap1 was the strongest, followed by Keap1α, whilst Keap1β exerted the weakest or least effects.

### 2.6 Distinct Keap1-based cellular responses of the Smad2/3 signalling to stimulation by TGFβ1, DTT or tBHQ

To examine distinct inducibly responsive effects of Keap1, Keap1α or Keap1β on the expression of Smad2/3 and cognate co-target genes, distinct genotypic cell lines *Keap1^+/+^*, *Keap1^−/−^*, *Keap1-restored*, *Keap1α-restored* and *Keap1β* (*Keap1^△1-31^*) were treated for 0 to 24 h with TGF-β1 (0.2ng/mL), DTT (1 mM, as a strong reducing agent(25)) or 50 μM of *tert*-butylhydroquinone (tBHQ, as a pro-oxidative stressor and also widely-used Nrf2 activator(37)). As expected, both the mRNA and protein expressed levels of all the examined Nrf2, Nrf1, Smad2, Smad3, Smad4, TGF-β1, E2F4, PAI-1 and TMEPAI were significantly augmented to certainly varying extents within distinct Keap1-based cellular responses to 24-h stimulation by TGF-β1 (Fig. 6), DTT (Fig. S8) or tBHQ (Fig. S9), when compared with those counterparts of each cell line treated at 0 h (i.e. basal control values).

**Figure 6.**
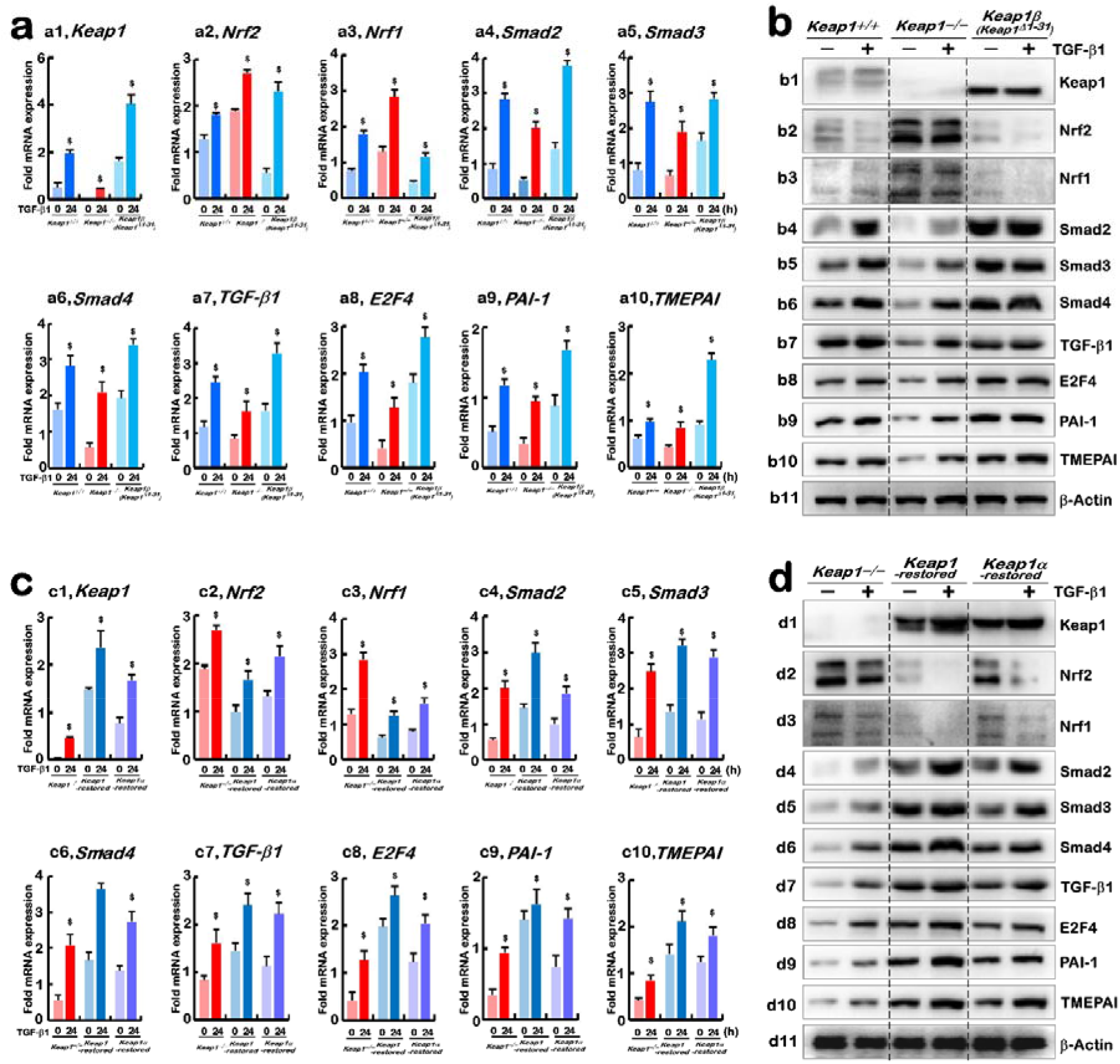
Distinction in DTT-stimulated expression of all examined genes in distinct Keap1-based genotypic cell lines. **a-b.** Each of experimental *Keap1^+/+^*, *Keap1^−/−^*, and *Keap1β(Keap1^Δ1-31^)* cell lines was (or was not) treated with 1 mM DTT for 0 or 24 h, before basal and DTT-inducible mRNA levels (*a*) and protein abundances (*b*) of Keap1, Nrf2, Nrf1, Smad2, Smad3, Smad4, TGF-β1, E2F4, PAI-1 and TMEPAI were determined by real-time qPCR and Western blotting, respectively. β -actin served as a loading control. **c-d.** Each of *Keap1^−/−^*, *Keap1-restored, Keap1α-restored* cell lines was (or was not) treated with 1 mM DTT, and then subjected to real-time qPCR (*c*) and Western blotting (*d*) as described above. Of note, all the above-described data as shown herein were representative of at least three independent experiments (n = 3 x 3), with significant decreases (*, *p* < 0.01) and significant increases ($, *p* < 0.01) being calculated, relative to the corresponding controls, respectively.

Such basally-expressed mRNA and protein levels of Smad2, Smad3, Smad4, TGF-β1, E2F4, PAI-1 and TMEPAI were strikingly downregulated by the loss of *Keap1^−/−^* (Fig. 6, a & b), but after treatment of *Keap1^−/−^* cells with TGF-β1 for 24-h, their stimulated expression abundances were modestly increased to certain degrees, when compared with their corresponding wild-type *Keap1^+/+^* controls. By contrast, the transcriptional expression of *Nrf2* and *Nrf1* was marginally induced by TGF-β1 (Fig. 6a), whereas both protein abundances appeared to be unaffected by this growth factor (Fig. 6b), albeit their basal expression levels were obviously enhanced in *Keap1^−/−^* cells. Conversely, *Nrf2* and *Nrf1* were basally down-regulated in *Keap1β* (*Keap1^△1-31^*) cells, but TGF-β1-stimulated mRNA expression of *Nrf2* was augmented, whereas *Nrf1* was only marginally stimulated by this growth factor in this Keap1β-expressing cells (Fig. 6a), but both protein abundances were further suppressed after treatment with TGF-β1 (Fig. 6b). In the meantime, basal and TGFβ1-stimulated expression levels of Keap1, Smad2, Smad3, Smad4, TGF-β1, E2F4, PAI-1 and TMEPAI were significantly elevated to different extents in Keap1β-expressing cells (Fig. 6, a & b). Collectively, these indicate that Keap1 exerts a positive regulatory effect on the TGFβ1-triggered Smad2/3 signalling response, but such effect of Keap1 appears to be independent of stimulation of Nrf2 and Nrf1 by this growth factor.

Similar positive effects of Keap1 on the TGFβ1-Smad2/3 signalling as described above were further fortified in both *Keap1-restored* and *Keap1α-restored* cell lines (Fig. 6, c & d), when compared to *Keap1^−/−^* cells. Basal and TGFβ1-induced expression of Keap1, Smad2, Smad3, Smad4, TGF-β1, E2F4, PAI-1 and TMEPAI at their mRNA and protein levels was all upregulated by restoration of Keap1 or Keap1α, but both protein, rather than mRNA, abundances of Nrf2 and Nrf1 were downregulated by TGFβ1 in *Keap1-restored* and *Keap1α-restored* cell lines, even though the basal protein expression of both CNC-bZIP factors was enhanced in *Keap1^−/−^* cells.

Next, the redox-inducible effects of Keap1 Keap1α or Keap1β on the TGFβ1-TGFSmad2/3 signalling responses to DTT or tBHQ were further determined herein (Figs. S8 & S9). Such redox-stimulated increases of all the examined gene products were observed in *Keap1^+/+^* cells, even in *Keap1^−/−^* cells they were also partially induced by DTT or tBHQ. Of note, basal and redox stimulated increases of Smad2, Smad3, Smad4, TGF-β1, E2F4, PAI-1 and TMEPAI were all detected in *Keap1β*(*Keap1^△1-31^*) cells, whereas Nrf2 and Nrf1 were augmented in *Keap1^−/−^* cells, as compared with their equivalent controls of *Keap1^+/+^* cells (Figs. S8 & S9). Conversely, restoration of Keap1 or Keap1α enabled such redox-stimulatory increases of Nrf2 and Nrf1 (with an exception of Keap1α stimulated by DTT) to be reduced, but stimulated increases of Smad2, Smad3, Smad4, TGF-β1, E2F4, PAI-1 and TMEPAI to be promoted, differentially in *Keap1-restored* and *Keap1α-restored* cell lines. Altogether, these demonstrate that redox-stimulated expression of Smad2/3 are also monitored by Keap1-dependent signalling and Keap1-indepdenpdent pathways. Overall, this study indicates that Keap1, together with its isoforms α and β, provides a novel functional crosstalk between the relevant redox signalling (e.g., mediated by Nrf2 and/or Nrf1) and its interactors Smad2/3 involved in the growth and development.

## 3. Discussion

In this study, we have identified that two Keap1 isoforms α and β are differentially expressed in distinct cell lines and also exerted discrepant effects on Nrf2 protein stability and transcriptional activity to regulate *ARE*-driven target genes (Fig. 7a). This finding just provides a novel explanation of why Nrf2, as a versatile chameleon-like regulator of antioxidant, detoxification, cytoprotective, and other genes in selective responses leading to repressing or promoting cancer (38), must have to be highly tightly regulated by a finely-tuned control system (involving Keap1 and its isoforms α and β with their direct and/or indirect interacting protein networks). Upon stimulation of cells by oxidative stress, chemopreventive compounds can block the activity of Keap1-Cul3-Rbx1 ubiquitin ligase (39), such that Nrf2 does not only circumvent the ubiquitin-mediated proteasomal degradation and is also precisely transactivated to properly regulate transcriptional expression of its downstream target genes. Thereby, it is inferable that the protein expression abundance of Nrf2 and its turnover after its functional action are all accurately fined tuned, to certain proper extents, by alternative translation of Keap1 into two isoforms α and β and by differential functioning of the ubiquitin-mediated proteasomal degradation system triggered by three distinct Keap1 dimers (αα, αβ, ββ). This is also modulated by its naturally-occurring dominant-negative mutant Keap1^△C^ lacking the most essential portion of the C-terminally Nrf2-interacting Kelch/DGR domain (40).

**Figure 7.**
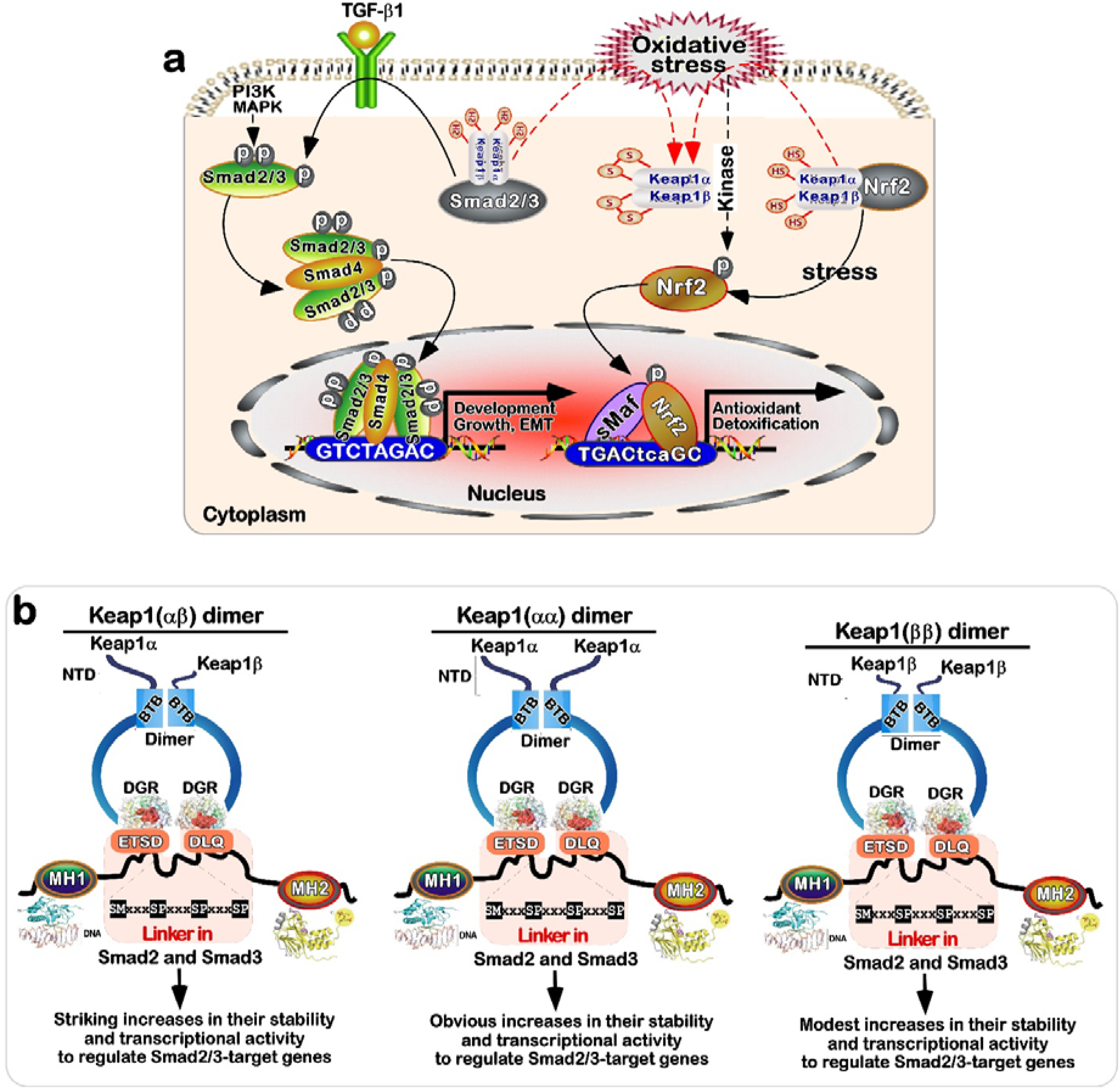
A novel model proposed for a newly-identified functional cross-talk of Keap1, as its isoforms α and β, with its interactors Nrf2 and Smad2/3. Of note, the stability of Smad2/3 and their transcriptional activity to mediate target gene expression are enhanced by physical interaction of Keap1 with two highly conserved EDGETSD and DLQ motifs within the linker region of Smad2/3 (*b*), although Nrf2 is negatively regulated by Keap1 in a similarly-binding fashion (*a*). Overall, this study presents a novel functional bridge of Keap1, Keap1α or Keap1β crossing both the redox-responsive Nrf2 signalling and the developmental TGFβ1-Smad2/3 pathways. In addition, this is also further supported by our experimental evidence revealing that the phosphorylated Smad2 activation occurs *via* MAPKs and PI-3 kinase signalling pathways in response to TGF-β1 and UVA (as oxidative stress) (Fig. S10).

Considering the strict specificity of the relationship between Keap1 and Nrf2 (41-43), it is hence hypothesized that the Keap1 isoforms α and β can exert their distinct intrinsic inhibitory effects on Nrf2 by physical interaction with the latter ETGE and DLG motifs to target this CNC-bZIP protein to the ubiquitin-mediated proteasomal degradation pathway. Keap1α is the full-length prototypical Keap1, whereas Keap1β arises from the secondary internal translation start codon to yield an N-terminally 31aa-truncated mutant Keap1^△1-31^. Such nuanced two Keap1 isoforms α and β have disparate half-lives, which are determined to be significantly shortened (from 9.4 h to 6.4 h) or prolonged (from 9.4 h to 28.9 h), respectively. It should also be noted that 9.4 h is the half-life of ectopic Keap1, but that of endogenous Keap1 is only 2.1 h). From these, taken together with our confocal imaging and subcellular fractionation data, it is thus reasoned that the three putative functional dimers of Keap1 (αα, αβ, and ββ) are manifested with its interactors (e.g., Nrf2, p62, Smad2/3, and others (32), and that they enable to distribute tempo-spatially in distinct subcellular (i.e., cytosolic, mitochondrial and nuclear) locations and hence exert distinctive regulatory effects and relevant biological functions. Yet, it warrants further study of the mechanistic distinction in the subcellular compartmentation of Keap1α and Keap1β to be tethered.

As expected, several lines of experimental evidence have been provided revealing that Keap1, Keap1α and Keap1β have exerted similar, but yet different, inhibitory effects on those examined Nrf2-target genes, although they all enable to physically interact with this CNC-bZIP factor(24). Such differential expression genes (DEGs) monitored by Keap1, Keap1α and Keap1β are scrutinized by the transcriptome sequencing of distinct Keap1-based genotypic cell lines (i.e., *Keap1^+/+^*, *Keap1^−/−^*, *Keap1-restored*, *Keap1α-restored* and *Keap1β*(*Keap1^△1-31^*)). The similarities and differences between Keap1, Keap1α and Keap1β in governing those gene expression profiles were further parsed by detailed KEGG pathway and GO enrichment analyses. Of striking note, a surprising association of Keap1 with Smad2 was, for the first time, found herein, and further validated by mass spectrometry analysis of Keap1-putdown interactome, implying there exists a competitive relevance of the Keap1-Nrf2 signalling with the TGFβ-Smad2/3 pathways (Fig. 7a). For instance, Nrf2 can inhibit the TGF-β1 expression of in a human stellate cell line (22), but conversely TGF-β1 can also enable downregulation of Nrf2 in another rat hepatic stellate cell line (23).

Subsequently, further experimental evidence has also been herein provided substantiating that Keap1, as well as its isoforms α and β, enables for a physical interaction with the two highly conserved EDGETSD and DLQ motifs of Smad2/3 (respectively compared with the ETGE and DLG motifs in Nrf2 enabling it to bind a dimeric Keap1 (44,45)), which are located within the central linker regions of Smad2/3 between the N-terminal MH1 (binding target genes) and its C-terminal MH2 (forming a functional heterodimer with Smad4, as illustrated in Figs. 7b). Notably, such an interaction of Keap1, Keap1α or Keap1β with Smad2 or Smad3 was corroborated by their co-immunoprecipitation and co-localization assays. These collective data demonstrate that the redox sensor Keap1 is likely involved in cross-talking with the Smad2/3-mediated signalling (Fig. 5a), although this pathway is triggered predominantly by the TGF-β family of growth factors(46). In this signalling cascade, TGF-β1 stimulates activation of Smad2/3 and in turn, Smad2/3 had also been shown to be central to the expression of TGF-β1 and Smad4 (as a co-binding partner of Smad2/3 recruited their cognate target genes, e.g., E2F4, PAI-1 and TMEPAI)(47), to comprise a positive feedback regulatory circuit.

Importantly, we have also found that the protein stability of Smad2/3 and its transcription activity are markedly enhanced to different extents by Keap1, Keap1α or Keap1β, although they enable both Nrf2 stability and *trans*-activity to be significantly reduced. This notion is supported by evidence revealing that the half-lives of endogenous Smad2 and Smad3 proteins are determined to be 1.4 h or 1.1 h, respectively, after CHX treatment, but ectopic Smad2 half-life is prolonged to 3.1 h, 2.2 h or 1.6 h, while ectopic Smad3 half-life is also extended to 2.4 h, 1.7 h or 1.3 h, by ectopically-expressing Keap1, Keap1α or Keap1β, respectively. Such opposing effects of Keap1, Keap1α or Keap1β on Smad2/3 and Nrf2 demonstrate a significantly mechanism accounting for stability of Smad2/3 and its turnover (Fig. 3d), which is much likely to be distinctive from the Nrf2-controlling mechanism, albeit t this remains to be further elucidated. Anyways, it is inferable that putative conformation changes in the Smad2/3’s functional complexes are likely to result from the physical interaction of dimeric Keap1 with two highly conserved EDGETSD and DLQ motifs in their central linker regions. This is due to the facts that this linker region includes a transactivation domain(48), enables homo-or hetero-oligomerization and provides critical phosphorylation sites by GSK3β, MAPKs, CDKs and calcium-calmodulin dependent kinases, such that it can exert key regulatory roles in governing Smad2/3 activity and function(49,50). For example, the linker region phosphorylation cannot only regulate the inhibition of the nuclear translocation of Smad2/3, and also influences their protein degradation by the ubiquitin-proteasomal system (11,51). This is also further evidenced by our experiments showing that the linker region is essential for stability of Smad2/3, possibly monitored by Keap1-adapted deubiquitinating system, but the latter detailed mechanism is required for further studies.

In addition to its interactors Smad2/3, other 72 transcription-related factors (e.g., STAT1/3, YY1, FOXK1, CTCF, NKRF, RELA, JDP2 and TFAM) were directly or indirectly put down by Keap1 (Fig. 3d), implying that Keap1 plays a vital role in multiple gene regulatory networks controlled by such many transcription factors. Interestingly, Keap1-interactome analysis also unravelled that it is likely to act as a key player in the process of sequentially adding and removing ubiquitin by dynamic formation of distinctive functional complexes with either ubiquitin-conjugating enzymes (e.g., Cullin-1, 2, 3, 4B, 5, ATG3, UBE3A, and UBE3C) along with proteasomal subunits or other deubiquitinating enzymes (e.g., USP5, 7, 10, 12, 14, 15, 17L, 22, 24, 47 and UCHL5), as well with cullin-associated NEDD8-dissociated protein 1 (CAND1, as an inhibitor promoting dissociation of substrate receptor components from Cullin RING ligases(52)). For this, it is inferable that such selective dynamic formation of two opposite functional complexes is much more likely to depend on distinctive tempo-spatial contexts between Smad2/3 and Nrf2. Thereby, based on our experimental data, it is deduced that after inhibiting the expression of Nrf2, Keap1 activates Smad2/3 and promotes the expression of TGF-β1, Smad4 and relevant targets such as E2F4, PAI-1 and TMEPAI. This is further evidenced by our data obtained from *Nrf1α^−/−^* (with diminished Keap1 abundance) and *Nrf2^−/−^* (with enhanced Keap1 expression levels) hepatoma cells, and *Keap1^−/−^* MEFs. In the meantime, it is intriguing to note that such a table of Smad2/3-target genes was also induced by redox stimulation by DTT and tBHQ, particularly in *Keap1β*(*Keap1^△1-31^*), *Keap1-restored* and *Keap1α-restored* cell lines. In turn, distinctions in the Smad2/3 signalling activation by TGF-β1 were observed in different Keap1-based cell responses to this growth factor. Importantly, there may exist a coordinated mechanism between Nrf2 (and Nrf1) and Smad2/3 possibly through Keap1 or its isoforms.

In summary, we have herein discovered two nuanced Keap1 isoforms α and β that arise from alternative translation starting sites within its mRNA transcripts and also interact with Smad2/3, besides Nrf2. Their similarities and differences in monitoring relevant gene expression networks were further interrogated by transcriptome sequencing and Keap1-interactome analysis, along with the routine reductionist experiments. Subsequent series of evidence unravelled distinct and even opposing effects of Keap1, as well as its isoforms α and β, on Nrf1 and Smad2/3 at their protein stability and transcriptional activity to mediate distinct sets of target genes. Of note, Keap1 was also uncovered to monitor relevant protein stability or turnover by selectively interacting with either ubiquitin-conjugating enzymes or deubiquitinating enzymes so as to form dynamic distinct functional complexes. Overall, this study presents a novel functional bridge of Keap1, Keap1α or Keap1β crossing both the redox-responsive Nrf2 and the developmental TGFβ1-Smad2/3 signalling pathways. Thereby, this discovery also provides a novel helpful understanding of the ‘double-edged sword’ effects of Keap1-Nrf2 or TGFβ1-Smad2/3 signalling on paradoxically suppressing or promoting cancer and other diseases.

## 4. Materials and Methods

### 4.1 Chemicals and antibodies

All chemicals were of the best quality commercially available. Cycloheximide (CHX) and TGF-β1 (transforming growth factor-β1) was purchased from Sigma⍰Aldrich (St Louis, MO, USA), whereas DTT and tBHQ were from MedChemExpress (MCE). The antibody against Nrf1 proteins was saved in our labortory (as previously described(53)). Anti-V5 and Anti-Flag monoclonal antibodies were from Invitrogen, with all other primary antibodies from Abcam. The β-actin antibody, as well as the secondary antibodies, were from ZSGB-BIO (Beijing, China).

### 4.2. Cell lines and transfection

Four distinct genotypic cell lines (*Kepa1^−/−^, Keap1-Restored, Keap1α-Restored* and *Keap1β(Keap1^△1-31^)*) were created from wild-type HepG2 cells (i.e., *Keap1^+/+^*) described elsewhere(24). They were all allowed to grow in DMEM supplemented with 10% (v/v) foetal bovine serum (FBS), 5 mM glutamine, and 100 units/mL penicillin and streptomycin at 37 °C in an incubator with 5% CO_2_. The cells were transfected for 8 h with indicated plasmids in Lipofectamine 3000 (Invitrogen) and then allowed for 24-h recovery from transfection by changing a fresh medium before subsequent experiments.

### 4.3. Expression Constructs

Those expression constructs for human Keap1, Nrf2, CTTN, Smad2 and Smad3 were made by cloning each of their full-length cDNA sequences into a pcDNA3 or 3xFlag vector. The N-terminal 32^nd^ methionine of the full-length Keap1 was mutated (or deleted) to yield only Keap1α-expressing plasmids, whereas the first N-terminal 31 amino acids of the full-length Keap1 were deleted to yield the Keap1β plasmid. The other plasmids specifically for the genome editing of Keap1, and Smad2/3 mutants were made and identified (as shown in Figs. 1d-h & 4g). In addition, an *E2F4*-promoter luciferase reporter was also created herein. All plasmids were confirmed by sequencing and validated by relevant experiments.

### 4.4. Construction of *Keap1^−/−^*, *Keap1β*(*Keap1^△1-31^*), *Keap1-Restored* and *Keap1α-Restored* cell lines

Both Keap1 (sgRNA-Keap1: 5’-TATGAGCCAGAGCGGGATG-3’) and Keap1α (sgRNA-Keap1α: 5’-AGGCCTAGCGGGGCTGGG GC-3’) genomes had been created *via* CRISPR/Cas9 and identified previously (24). Then, each of those Cas9 plasmids for Keap1 and Keap1α were introduced into HepG2 cells to construct *Keap1^−/−^* and *Keap1β(Keap1^△1-31^)* cell lines (Fig. S1c). Thereafter, the full-length cDNA sequences of the Keap1 plasmid and Keap1α plasmid (F: CGCGGATCCATGCAGCCAGAT CCCAGG, R: AAATATGCGGCCGCTCATAGCCTCCTCTCCACACT) were subcloned into the pLVX-IRES-ZsGreen-puro vector for lentivirus packaging, which was verified by gene sequencing (Fig. S1d). These two viruses were prepared in 293T cells with three plasmids (2.52 μg pMD2G, 8.34 μg psPAX2, and 10.5 μg pLVX-mcmv-ZsGreen-puro pKeap1/pKeap1α) according to a previously-described method(54). Either of the Keap1 and Keap1α viruses was infected into *Keap1^−/−^* cells, and then screened in puromycin-containing media to select the positively-infected cells. The Keap1-or Keap1α-restored cells were verified by Western blotting.

### 4.5. RNA isolation and quantitative real-time PCR (qPCR)

Experimental cells were subjected to total RNA isolation (using an RNA Simple Total RNA Kit, DP419, Tiangen Biotech Co., Ltd., Beijing, China). The first strand of cDNA was obtained by adding total RNA (1.5 µg) in a reverse transcriptase reaction (using a Revert Aid First Strand cDNA Synthesis Kit, from Thermo), and served as the template for quantitative PCR (using GoTaq qPCR Master Mix, A6002, Promega, USA). Then, each pair of those reciprocal primers (listed in Table 1) was also added to the indicated qPCR, which was carried out under the following conditions: 95 °C for 5 min, followed by 40 cycles of 15 s at 95 °C and 30 s at 60 °C. The mRNA expression level of β-actin was used as the optimal internal standard control.

**Table 1.**
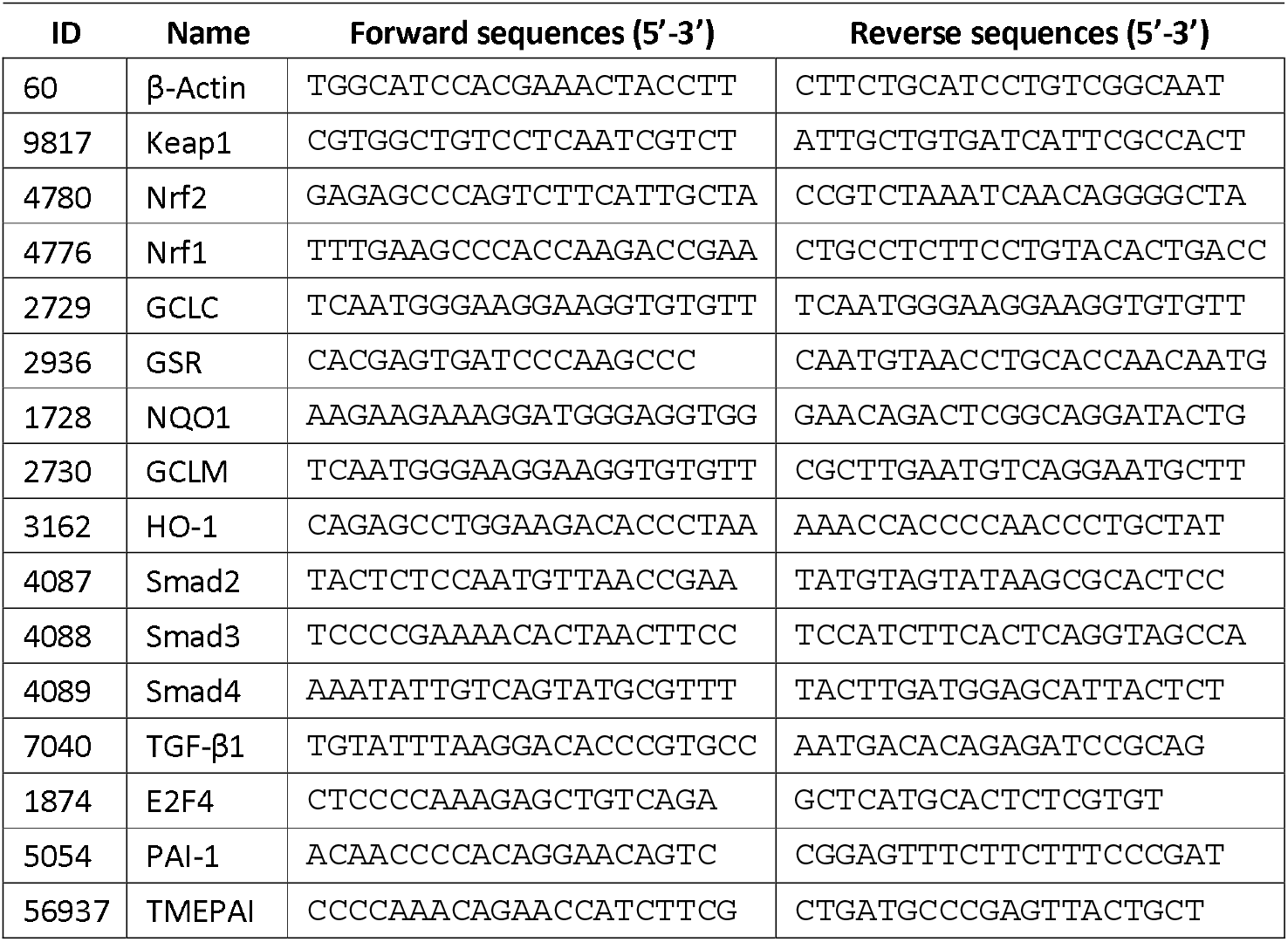
The pairs of used primers for the RT⍰qPCR analysis.

### 4.6. Luciferase Reporter Assay

Equal numbers (1.5 × 10^5^) of experimental cells were seeded in each well of 12-well plates. After reaching 70-80% confluence, the cells were transfected with luciferase plasmids alone or plus other expression plasmids in the Lipo3000 reagent, in which the *pRL-TK* plasmid served as an internal control for transfection efficiency. The luciferase activity was determined by the Dual-Lumi™ double luciferase reporter gene assay (RG088M, Beyotime).

### 4.7. Western Blotting analysis

After experimental cells were rinsed three times with PBS and harvested in a total protein extraction buffer, the total lysates were denatured immediately for 10 min at 100 °C, and then the eluted proteins were subjected to separation by SDS⍰PAGE. The indicated protein abundances were visualized by Western blotting with different primary and secondary antibodies.

### 4.8. Subcellular fractionation

Experimental cells (6 × 10^5^) were allowed for growth in 6-cm cell culture plates for 48 h and subjected to isolation of the cytoplasmic and nuclear fractions by using a relevant separation kit (N-3408-200ML, Sigma). During this procedure, the extracted cytoplasmic portion was further centrifuged again at 20,000 g for 10 min to obtain a precipitated mitochondrial fraction, while the supernatant was viewed as a cytosolic fraction. All these subcellular fractions were further evaluated by Western blotting with specific antibodies against indicated proteins and relevant compartmentally-localized markers

### 4.9. Transcriptome sequencing and protein profiling

Experimental cells were subjected to total RNA isolation using an RNA Simple Total RNA Kit, followed by transcriptome sequencing (Beijing Genomics Institute (BGI), Shenzhen, China) to obtain the relevant data from an Illumina HiSeq 2000 sequencing system (Illumina, San Diego, CA). All differentially expressed genes (DEGs) were identified. The standard fold change was ≥2 or ≤0.5, with FDR (false discovery rate) ≤0.001 determined by the Poisson distribution model method (PossionDis). The total proteins of COS-1 cells, that had been transfected with expression constructs for Keap1, Keap1α or Keap1β, were extracted and put down by Keap1-specific antibodies, followed by mass spectrometry analysis (BGI) to obtain those Keap1-interactome profiling data.

### 4.10. The CABS-dock structural modelling

The CABS-dock web server (http://biocomp.chem.uw.edu.pl/CABSdock/) was here used for the flexible protein-peptide docking, which enables full flexibility of the peptide structure with large-scale flexibility of the protein during the search for their binding site. The two putative KEAP1-binding peptides within SMAD2 (YISEDGETSD and NHSLDLQPV) or SMAD3 (YLSEDGETSD and HNNLDLQPV) were selected for docking into KEAP1 (PDB code: 1ZGK) within default settings. The ten top-ranked CABS-dock models were analysed.

### 4.11. Co-immunoprecipitation analysis

COS-1 cells (4.0 × 10^5^), that were grown on 6-well cell culture plates for 24 h, were transfected with different expression plasmids for Keap1-V5, Keap1α-V5, Keap1β-V5, Smad2-Flag or Smad3-Flag, and then allowed for 24-h recovery in the normal medium. The cells were lysed in RIPA buffer (C1053, PPLYGEN) containing protease and phosphatase inhibitors (Solarbio, P6730). The total lysates were centrifuged to collect the supernatants, followed by co-immunoprecipitation with antibodies against V5 or FLAG epitopes and the BeyoMag™ Protein A+G beads(55) (Beyotime. P2108-1ML). The eluted proteins were determined by Western blotting with anti-V5 and anti-FLAG antibodies (Invitrogen) and visualized by the horseradish peroxidase-conjugated secondary antibodies (ZSGB-BIO, ZB-2305), along with being developed by enhanced chemiluminescence reagents(56) (Thermo, iBright 750). Of note, an additional co-immunoprecipitation assay with antibodies against endogenous Keap1 (consisting of its α and β isoforms), followed by immunoblotting with antibodies against endogenous Smad2 or Smad3, was carried out in MHCC97L and THP-1 cell lines.

### 4.12. Immunocytochemistry and confocal microscopy

COS-1 cells (2.5 × 10^5^) were grown for 24 h on 6-well cell culture plates (with the glass coverslips of 1 cm^2^ c being stored flat) and co-transfected for 8 h with expression constructs for Smad2/3-Flag plus Keap1-V5, Keap1α-V5 or Keap1β-V5. The cells were allowed for 24-h recovery, then washed in PBS and fixed with paraformaldehyde fixative (Servicebio) for 30 min at room temperature. After the fixed cells were rinsed three times with PBS, they were permeabilized with 0.2% Triton X-100 for 20 min, before immunocytochemical staining. The samples were further rinsed again three times with PBS, blocked with 1% BSA for 60 min, incubated overnight with each of indicated primary antibodies and then re-incubated for 4 h with the secondary Alexa Fluor–conjugated secondary antibody (ZSGB-BIO). The antibodies-stained coverslips were mounted with DAPI (Beyotime, C1005), and subjected to confocal imaging by using an IN Cell Analyser ZeissLSM900 cellular imaging system(31).

### 4.13. Statistical analysis

The statistical significances of differential gene expression levels measured by real-time qPCR and luciferase activity were determined using Student’s *t*-test or two-way ANOVA. All the relevant data were obtained from at least three independent experiments and shown as fold changes (mean ± SEM or SD) with significant statistical differences being calculated by the value of *p* < 0.05 or 0.01 when compared with controls).

## Author Contributions

Both F.C. and M.X. performed the experiments with help of R.W. and S.H., collected all the relevant data, and wrote a draft of this manuscript with most figures and supplemental information. Q.W. done a number of bioinformatics analysis of both transcriptome sequencing and Keap1-interactome protein profiling. D.L. modelled the CABS-dock structural interaction of Keap1 with Smad2/3. Y.W. and Z.Z. provided invaluable and critical discussion. Z.Z. also edited and corrected this paper in the standard English language. Lastly, Y.Z. designed and supervised this study, analysed all the data, helped to prepare all figures with cartoons, rewrote and revised the paper. All these co-authors have read and agreed to the published version of the manuscript.

## Supplementary Materials

The supporting information, including thirteen supplemental figures and also eight supplemental tables.

## Declaration of interests

The authors declare no competing interests.

## Acknowledgments

We are greatly thankful to all the other present and past members of Zhang’s laboratory for giving critical discussion and invaluable help with this work. This study was funded by the National Natural Science Foundation of China (NSFC, with two projects 82073079 and 81872336) awarded to Prof. Yiguo Zhang (at Chongqing University). This is also, at part, supported by the Initiative Foundation of Jiangjin Hospital affiliated to Chongqing University (2022qdjfxm001).

## Conflicts of Interest

The authors declare no conflict of interest. Besides, it should be noted that the preprinted version of this paper had been initially posted at doi: https://doi.org/10.1101/2022.11.22.517594.

## Notes

### Competing Interest Statement

The authors have declared no competing interest.

### Summary of Updates

Some content has been revised.

